# Structural dynamics of Zika Virus NS1 via a reductionist approach reveal the disordered nature of its beta roll domain in isolation

**DOI:** 10.1101/2022.04.16.488568

**Authors:** Shivani Krishna Kapuganti, Prateek Kumar, Rajanish Giri

## Abstract

Flavivirus Non-structural 1 (NS1) protein performs multiple functions such as host immune evasion, interaction with complement system factors, membrane rearrangement, etc. Therefore, it is highly plausible that significant structural and folding dynamics of NS1 might play a role in its multifunctionality. The dimeric structures of NS1 of multiple flaviviruses, including Zika virus (ZIKV), are available. However, its domain-wise dynamics perspective has not been explored so far. Therefore, it is of utmost importance to understand the structural conformations of NS1 and its domains in isolation, possibly highlighting the implications on the overall NS1 protein dynamics. Here, we have employed extensively long molecular dynamic (MD) simulations to understand the role of monomer, dimer, and a reductionist approach in understanding the dynamics of the three structural domains (i.e., β- roll, wing, and β-ladder) in isolation. Further, we experimentally validated our findings using CD spectroscopy and confirmed the intrinsically disordered behavior of NS1 β-roll in isolation and lipid mimetic environments. We also found that the β-ladder domain is highly flexible during long simulations. Therefore, we believe this study may have implications for significant dynamics played by NS1 protein, specifically during oligomerization of NS1.

**Graphical Abstract:** 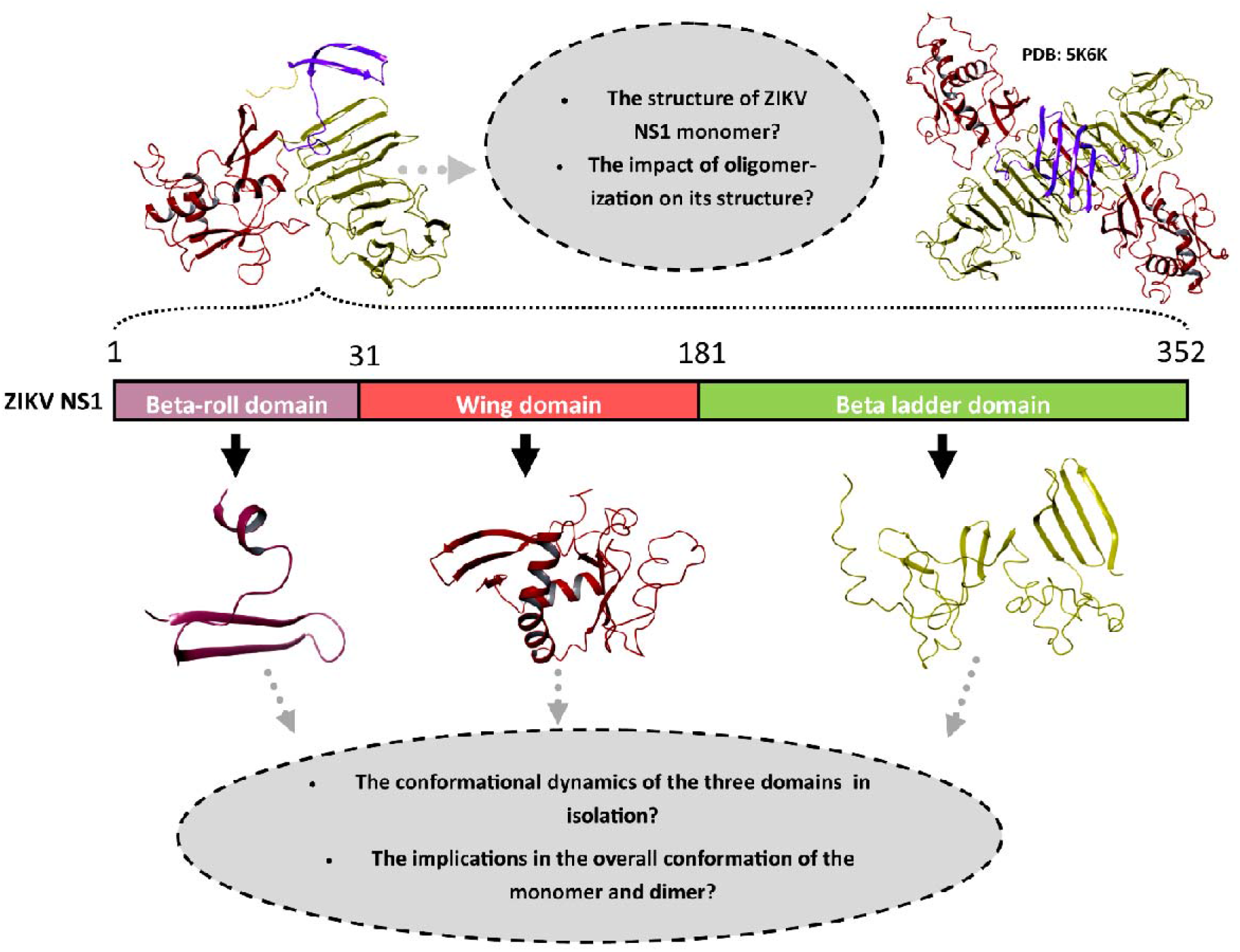

Schematic representation of the ZIKV NS1 protein and the models that we have used in this study.

## Introduction

ZIKV NS1 has multiple roles from immune evasion, cell surface interaction, and anchoring viral replication complex to autophagy.^1^ NS1 is capable of eliciting antibody response in the host. Both the viral protein and the elicited antibody seem to be responsible for the pathogenesis of ZIKV.^2,3^ NS1 is the only secreted flaviviral protein that may interact with a wide range of host proteins and may also have a role in recruiting them for aiding in viral replication.^4^ Since NS1 has been shown to interact with NS4A genetically; it has been hypothesized that NS1 may help anchor the replication complex.^5^

The ZIKV NS1 dimer crystal structure was elucidated in 2016 by Brown *et al*. As revealed in the structure, N-terminal amino acids, 1-30 form a β-roll, residues 31-180 form an epitope rich wing domain, and residues 181-352 form the C-terminal β-ladder domain. The cross-shaped dimer is formed via the intertwining of β-roll and end-to-end β-ladders. Regions lying within residues 30-37 and 152-180 of flavivirus NS1 form a subdomain that connects β-roll and β-ladder. This subdomain and β-roll together form a hydrophobic protrusion that contacts lipid membranes. Furthermore, β-roll is the most conserved region across different flaviviruses NS1 proteins.^6,7^

After translation, NS1 undergoes glycosylation and forms dimers, the functionally active form. A significant fraction is associated with the luminal side of viral replication vesicles.^8^ Some of it is transported to the Golgi, followed by glycoside trimming and incorporation of lipid moieties. NS1 is then secreted as a hexamer with a lipid core constructed by the association of three dimers together to form a trimer of dimers.^9,10^ They are held together by interacting with the same lipid raft composed of around 70 lipid molecules. The inner face of the dimer consisting of the β-roll and the connector subdomain interacts with the lipids.^6^ A report has demonstrated that lipid rafts disrupt compound, Methyl-B-cyclodextrin, and competitive inhibitors of the HMG-CoA reductase, a rate-limiting enzyme in cholesterol synthesis, lovastatin, and pravastatin significantly reduced secretion of DENV NS1 and viral load.^11^ Another study showed that knockdown or inhibition of cholesterol transporter complex CCC components resulted in a decrease in the secretion of DENV NS1 and viral load.^4^

In view of the multi-functionality of NS1, it is of utmost importance to understand the structural conformations of NS1 and its subdomains, mainly through reductionist approaches. Here, we employed MD simulations to understand the role of monomers and their association with the dimer structure. Then, we used a reductionist approach to understand better the dynamics of its three structural domains in isolation. Further, we tried to understand the folding of ZIKV NS1 β-roll domain in isolation, experimentally, in different lipid mimetic environments hoping that this would provide insight into its structural changes during interactions with different cellular membranes.

## Methodology

### Peptide and reagents

ZIKV NS1 β-roll domain (residues 1-30) (N-DVGCSVDFSKKETRCGTGVFIYNDVEAWRD-C) was synthesized and procured from ThermoFisher Scientific, USA. The purity of the peptide was >91%. The mass spectrum and the HPLC spectrum, and certificate of analysis of the synthesized peptide, have been attached in the supplementary file (**Figure S1**). The peptide was dissolved in PBS to prepare stock solution (0.25 mg/ml). The 2,2,2-trifluoroethanol (TFE) and Sodium dodecyl sulfate (SDS) were procured from Sigma-Aldrich (USA). The lipids 1,2-dioleoyl-sn-glycero-3-phosphocholine (DOPC), 1,2-dioleoyl-sn-glycero-3-phospho-L serine (DOPS), and 1,2-Dioleoyl-3-trimethylammonium propane (DOTAP) were obtained from Avanti Polar Lipids (Alabaster, Alabama, U.S.A.) and prepared as described in our previous studies.^12,13^

### Liposome preparation

The chloroform in the lipids was removed using rotary evaporators and further overnight incubation in a desiccator. The lipids were then subjected to five alternative cycles of freezing (in liquid nitrogen) and thawing (using 60_lJ_C water bath). The lipids were then diluted in water and extruded using a 0.1 μm pore diameter polycarbonate membrane and the Avanti mini-extruder (Avanti). The hydrodynamic radius/size of thus prepared LUVs (1:100 dilution in water) was measured using DLS (Zetasizer Nano S from Malvern Instruments Ltd., UK). The observed size of the LUVs was ∼100nm.

### Circular dichroism (CD) spectroscopy

J-1500 spectrophotometer (Jasco) was used to record all CD spectra. 1 mm length quartz cuvette was used to record the spectra at 25°C in various concentrations of SDS, LUVs, and TFE. 40 μM ZIKV NS1 β-roll domain peptide was dissolved in 1X PBS. The spectra were fitted with FFT filter smoothing (7 points of smoothing window).

### Lifetime analysis

The fluorescence lifetime of Trp in the β-roll domain was measured using the DeltaFlex TCSPC system (Horiba Scientific). The wavelengths for excitation monochromator was set up at 280 nm and emission monochromator at 350 nm.The measurement range was set up to 200 ns with 32 nm of bandpass and a peak preset of 10000 counts. Ludox was used as a prompt to correct the instrument response factor (IRF), and the wavelength for prompt measurement was set up at 280 nm. In addition, a 40 μM protein sample used for CD spectra was prepared in PBS and used for all measurements.

### 3D Structure modeling and Molecular dynamics simulations

For the dimer simulations, the crystal structure with PDB ID: 5K6K was used. For the monomer simulations, a single chain from the crystal structure of the dimer was used. The structure models for all three NS1 domains in isolation were built using the RaptorX webserver. The models were prepared by adding missing hydrogen atoms, improving improper bond orders, and parameterizing asymmetrical residues in Schrodinger’s protein preparation wizard using OPLS 2005 forcefield.

For MD Simulations of RaptorX generated structures, we have used our previously reported protocols.^14^ In brief, all simulation systems were set up in TIP3P water model and neutralizing ions in Gromacs v5 using Charmm36 force field. The average temperature and pressure were maintained at 300K and 1 bar, respectively, using Nose-Hoover and Parrinello-Rahman coupling methods during simulation. Further, to investigate the structural conformations of the β-roll domain in the presence of lipids, the simulation setups were prepared using CHARMM-GUI webserver. For this purpose, the DOPC and DOPS lipid molecules were used to surround the β-roll domain. Finally, the simulation trajectory analysis and calculations were performed using Maestro, Chimera, and Gromacs commands for calculating root mean square deviation (RMSD), fluctuation (RMSF), and protein structure compactness using radius of gyration (RG).

## Results and discussion

### 1. Secondary structure and disorder propensity prediction of ZIKV NS1 and its domains in isolation

To better understand the structural conformations of NS1 and isolated domains, PsiPred was used to predict the secondary structure, making predictions based on the intrinsic properties in the sequence. From PsiPred analysis, it was observed that full-length NS1 is predicted to contain 31.8% beta-strand structure, 47% alpha-helical structure, and 54.9% random coil. The isolated domains were predicted to have 50%, 50%, and 62.8% random coil in β-roll, wing, and β-ladder domains, respectively (**Figure S2**).

As previously reported by our group in 2016, the sequence-based disorder prediction of overall ZIKV NS1 protein has shown very less disordered regions.^15^ Here, we have checked the disorder propensity of its three domains in isolation, using multiple predictors with newly developed and improved algorithms. To overcome the biasedness of predictors, a set of five online predictors viz. PONDR-VLXT, PONDR-VL3-BA, PONDR-VSL2, IUPred2A, and PrDOS were used to predict the disorder propensity in ZIKV NS1 domains in isolation (**Figure 1**).

**Figure 1.**
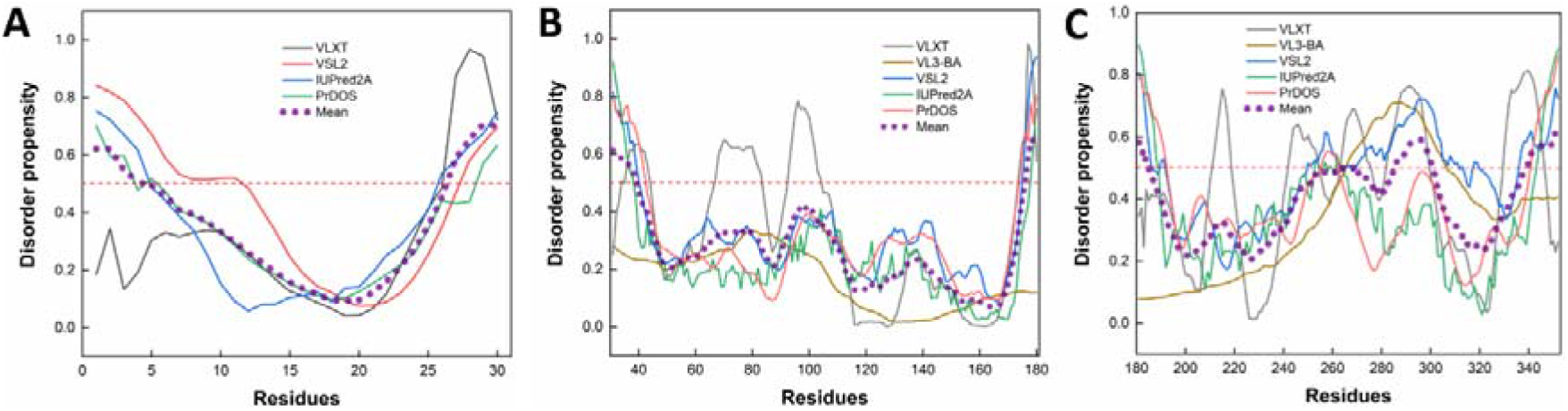
Disorder profiles generated using PONDR-VLXT, PONDR-VL3-BA, PONDR-VSL2, IUPred2A, and PrDOS online predictors of ZIKV NS1 domains in isolation (A) β-roll domain, (B) wing domain, and (C) ladder domain. The dotted line in purple represents the mean disorder propensity.

### 2. Molecular Dynamic Simulations of ZIKV NS1; dimer, full-length monomer and its domains in isolation

Elaborate MD simulations were performed to understand the dynamics of isolated domains. We also performed monomer and dimer simulations to compare our results of the isolated domains and investigate in the perspective of monomer and dimer dynamics. The results of monomer and dimer simulations correlate well with the available literature on the same.^16,17^ The 3D structure of the full-length ZIKV NS1 dimer was retrieved from Protein Data Bank (PDB ID: 5K6K), and missing residues were modeled using Prime module in Schrodinger. In isolation, the three domains of NS1 (β-ladder, wing and β-roll domains) were modeled using RaptorX. The MD simulations were done on the RaptorX generated models using GROMACS software.

#### 2.1 MD simulations of ZIKV NS1 dimer

The NS1 homodimer is the functionally active form. Two NS1 monomers interact to form a cross-shaped dimer, where the β-roll domains of two monomers intertwine with each other. Similarly, the β-ladders of two monomers interact with each other in an end-to-end manner. Owing this arrangement, the dimer has a hydrophobic inner face and a polar outer face. On the inner face, the β-roll domain, greasy finger, and the connector subdomains come together and form a hydrophobic surface.^18^ The rest of the protein arranges into the polar outer face. A trimer of dimers comes together to form the hexamer lipoprotein secreted out of the cells.^9^

In **Figure 2B**, the structures of the dimer before (yellow) and after simulation (blue) have been superimposed. Upon simulations of the dimer up to 100 ns, the rigidity of the dimer is maintained and fluctuates less between 3.04 - 3.14 nm **(Figure 2C)**. Likewise, the RMSD values, even after reaching the plateau phase, stay below 0.34 nm **(Figure 2D)**. Further, the RMSF values of both the dimer chains do not show significant fluctuations at the N-terminal region, corresponding to the β-roll domain, contrary to what was observed in the monomer simulations (**Figure 2E**; inset shown in black outline). However, a specific region (residues ∼100-130) of the wing domain, which is implicated in the interaction of NS1 with various cellular factors, showed high fluctuation in RMSF values in the dimer simulations (**Figure 2E**; inset shown in yellow outline).

**Figure 2.**
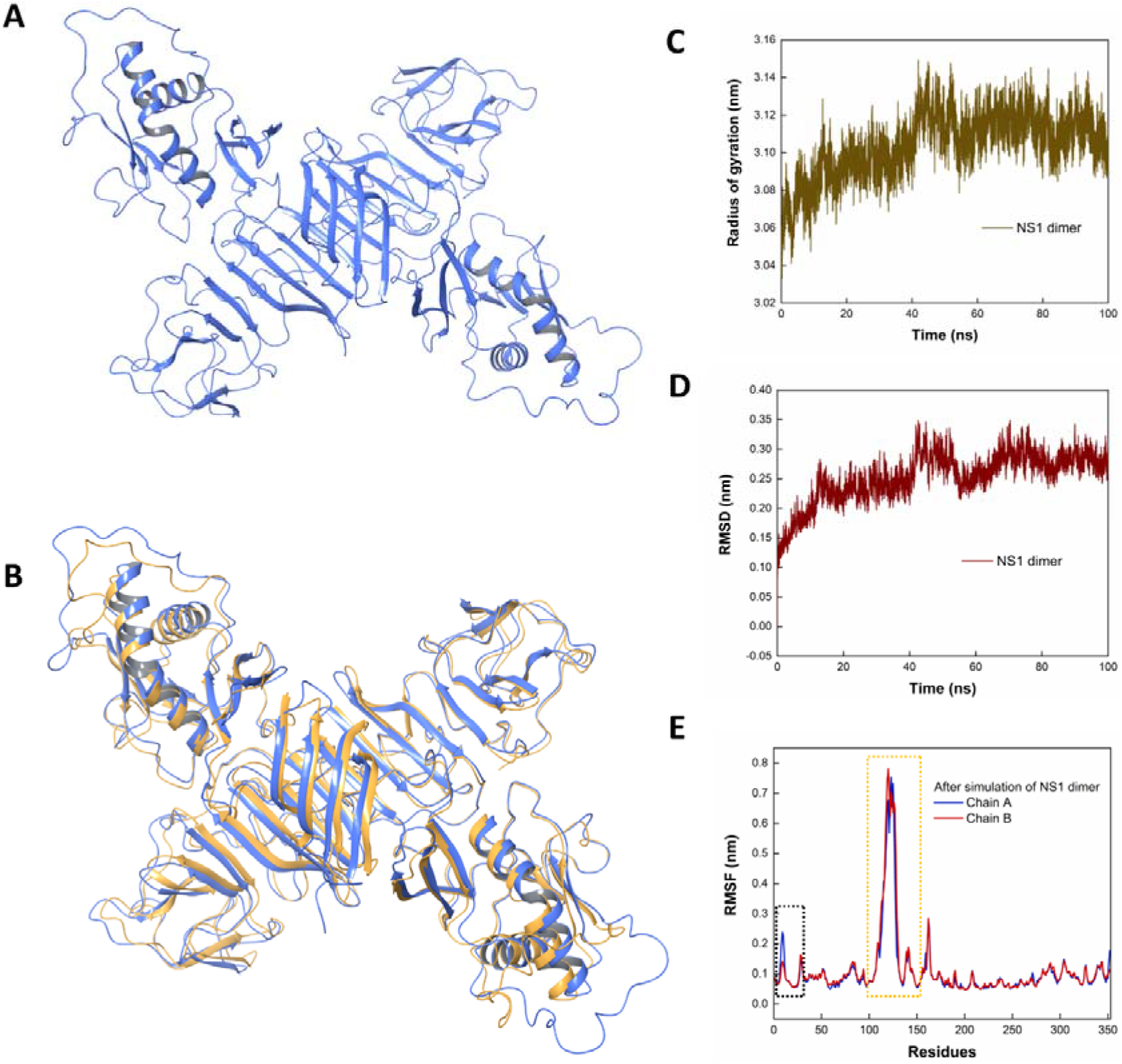
Molecular Dynamic simulations of (A) ZIKV NS1 dimer in native water environment and (B) superpositioned 100 ns frame over the crystal structure (PDB ID: 5K6K), yellow represents the structure before simulation and blue after simulation. (C), (D), and (E) show the graphical representations of Radius of gyration, RMSD and RMSF, respectively, on simulation of ZIKV NS1 dimer in water up to 100 ns. Insets in (E) show the residue positions of β-roll and wing domains. .^6, 7^

#### 2.2 MD simulations of ZIKV NS1 monomer

The full-length ZIKV NS1 has three structural domains; the β-roll domain at the N-terminal, central wing domain, and C-terminal β-ladder domain. It has 12 cysteine residues that form six disulphide linkages within the monomer. Compared to WNV and DENV NS1 proteins, the ZIKV NS1 has an extended hydrophobic surface due to the combined effect of the β-roll, greasy finger, wing flexible loop, and the connector sub-domains.^6,7^

The structure of full-length ZIKV NS1 monomer had a mixed α-β structure. However, a few structural changes were observed after 100 ns simulation in water, as shown in **Figure 3A**. In Figure 3B, the monomer structures before (red) and after simulation (purple) have been superimposed and clearly illustrate their differences. In particular, the N-terminal region corresponding to β-roll domain (residues 1-30) has shown high fluctuations and loosening of structure after simulation. Due to the structural transitions at N-terminal and other positions, the compactness of the structure is varied (from 2.30 nm to 2.42 nm) during initial simulation time (**Figure 3C**). The RMSD values stabilize after 60 ns of simulations but have a value higher than 0.4 nm, indicating that the monomer can undergo dynamic conformational changes (**Figure 3D**). From the RMSF values, we can further localise the conformationally dynamic regions to the N-terminal region, which comprises the β-roll domain (**Figure 3E**; inset shown in black outline) and certain regions of the wing domain (**Figure 3E**; inset shown in yellow outline). The behaviour of the N-terminal region is different in the dimer and the monomer. This may indicate that the interaction of the β-roll domain, which is involved in oligomerization, with the β-roll domain of the other chain, restricts the conformational flexibility of this region.

**Figure 3.**
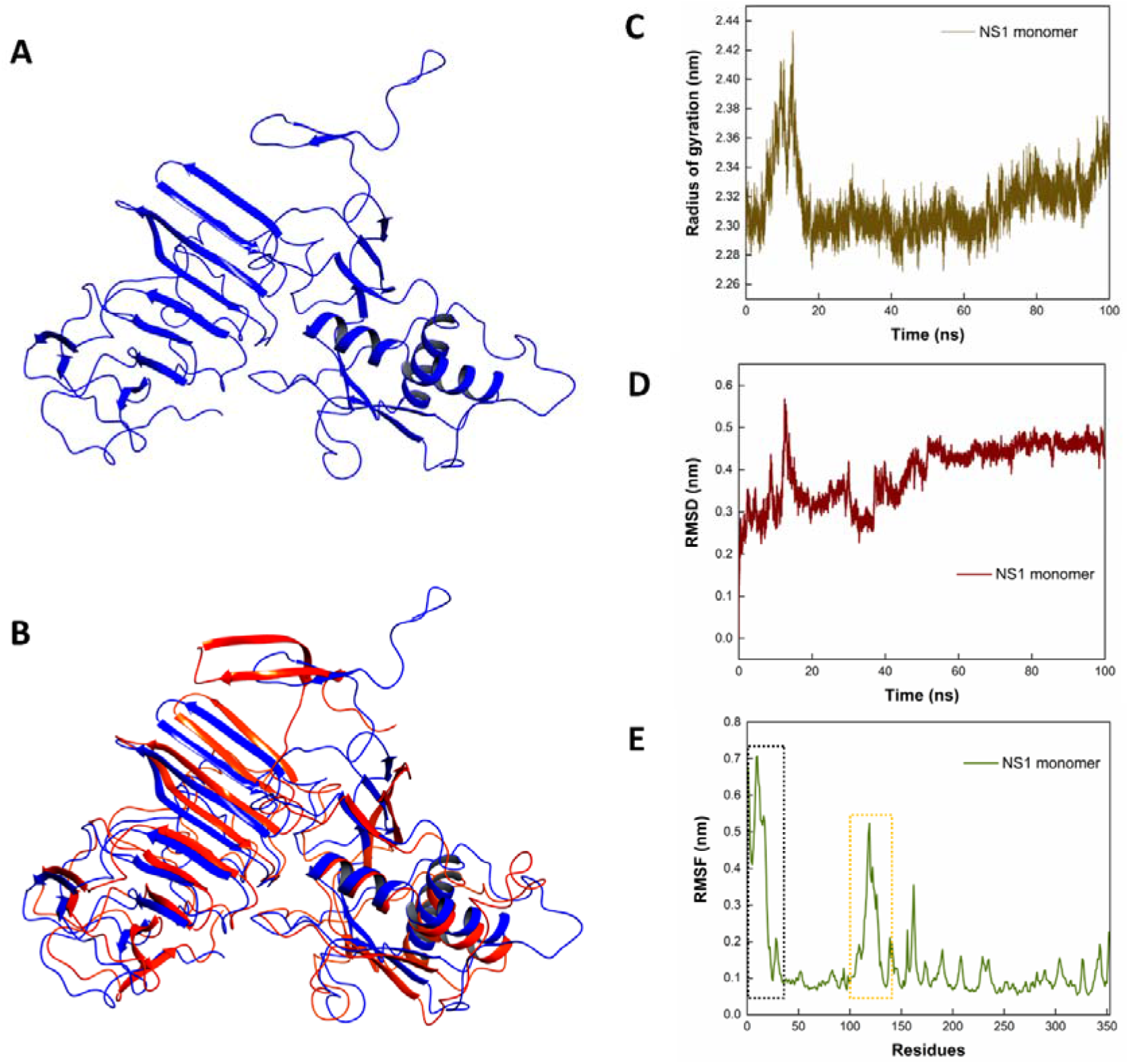
Molecular Dynamic simulations of (A) full-length ZIKV NS1 monomer in native water environment. (B) shows the superpositioned 100 ns frame, in blue, over the structure of the single-chain from crystal structure (PDB ID: 5K6K), shown in red. (C), (D) and (E) show the graphical representations of Radius of gyration, RMSD, and RMSF, respectively, on simulation of ZIKV NS1 monomer in water up to 100 ns. Insets in (E) show the residue positions of β-roll and wing domains

Two groups have shown the MD simulations of ZIKV NS1, earlier.^16,17^ Our observations on the monomer and dimer simulations confirm the literature. Further, in order to get an overall perspective, we have conducted an unbiased study by taking a reductionist approach. The simulations of the three functional domains of ZIKV NS1 were also done to study their structural conformations in native water environment. ZIKV NS1 C-terminal β-ladder, central wing and N-terminal β-roll domains’ structures were generated on RaptorX and prepared for simulations as described earlier. The C-terminal β-ladder, central wing and N-terminal β-roll domains were simulated up to 500 ns.

#### 2.3 MD simulations of ZIKV NS1 β-ladder

The C-terminal β-ladder domain is the predominant structural feature of the NS1 dimer as it acts as the central axis around which the dimer stabilises.^6^ β-ladder domains of two monomers join together in an end-to-end manner and 20 beta strands, 10 from each monomer, arrange themselves together like rungs on a ladder. Most of the loops between the strands are short except for a long *spaghetti* loop between strands 13 and 14. The β-ladder domain contains the second glycosylation site at residue N207. The beta sheets of the β- ladder domain together with the β-roll domain and the second half of intertwined loop of the wing domain form a hydrophobic plane, and on the other side, the loops of the β-ladder domain including the spaghetti loop, its C-terminal end, the first half part of intertwined loop of the wing domain and its N130-glycosylation site form a hydrophilic plane.^7^

Here, in this study, we have modelled a structure of β-ladder domain of NS1 protein of ZIKV. As per the structure model, a mixed population of beta strands and unstructured regions are seen. **Figure 4A** shows the simulations of β-ladder domain in isolation up to 500 ns. The radius of gyration, RMSD and RMSF values have been plotted after 500 ns of simulations (**Figure 4B-4D**). The RMSD values reach a plateau phase above 1.2 nm which indicates a significant level of structural dynamics (**Figure 4C)**. Residues responsible for fluctuations in the RMSF values are mostly contributed by residues involved in forming loops between individual beta strands. (**Figure 4D)**. This indicates that these regions of the beta ladder domain may exhibit dynamic conformations in different environments and upon interaction with different binding partners. Also, the C-terminal region, which is fused to NS2A, where proteolytic cleavage occurs also shows high fluctuations in RMSF values.

**Figure 4.**
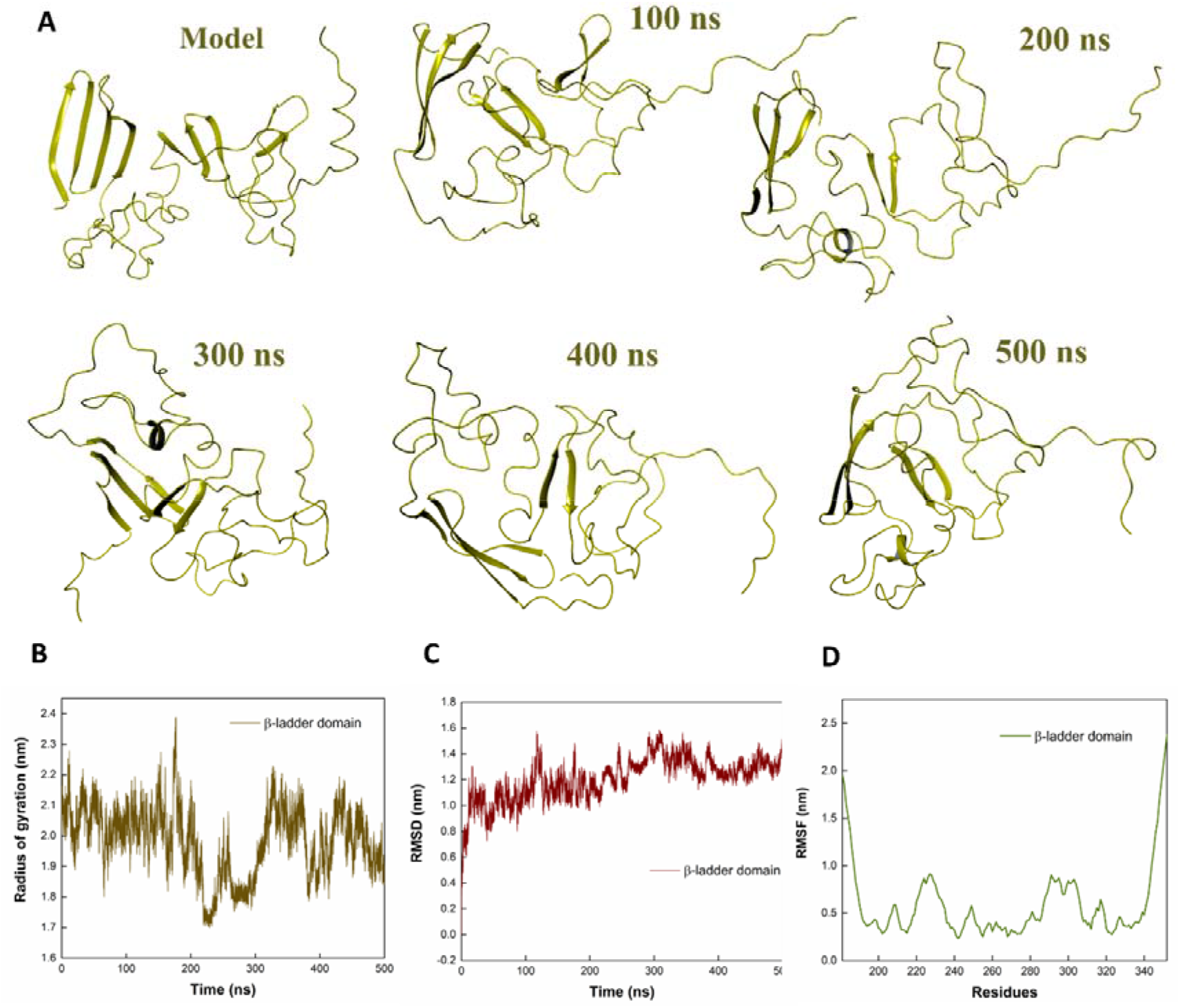
Molecular Dynamic simulations of ZIKV NS1 β-ladder domain in isolation in water up to 500 ns (A) ZIKV NS1 β-ladder domain in isolation in native water environment at 100, 200, 300, 400 and 500 ns. (B), (C), and (D) show the graphical representations of Radius of gyration, RMSD and RMSF, respectively. The comparatively high RMSF values observed corresponds to the loop regions that connect beta strands.

#### 2.4 MD simulations of ZIKV NS1 wing domain

The central wing domain sticks out of the cross shaped dimer structure, akin to the appearance of a pair of wings, hence, the name. It has a glycosylation site at N130. This domain has tremendous functional role in interaction of NS1 with various cellular factors. This region is polar in nature making it a good candidate for interactions in polar solvents.^7^ It is also called the epitope rich wing domain as it has been implicated in interaction with factors of the immune system. NS1 is known to have immune system evasion properties, activating complement system and cross reacting with anti-bodies. In ZIKV NS1, of the Ugandan strain, the residues 108-129 form a wing domain flexible loop with properties that are not observed in WNV or DENV NS1 structures.^7^ This region significantly contributes to the hydrophobicity of the inner face of the dimer, along with the β-roll domain and the greasy finger residues 162-164. In the crystal structure of NS1 elucidated by Brown et al, they show that the residues W28, W115, W118 and F123 protrude out of the hydrophobic surface of the protein, therefore, may be involved in membrane interaction.^7^ The involvement of W115 in interaction of NS1 with envelope protein has been reported in DENV.^19^ In the crystal structure of ZIKV NS1 of Brazilian strain, elucidated by Xu *et al*, they show that the long loop (residues 91-130) harbours the hydrophobic spike made of residues Y122, F123 and V124.^18^ Scaturro *et al* show that probably, this spike is essential for NS1 interaction with envelope protein in DENV.^19^

Similar to the experimentally determined structure, the wing domain model in isolation contain a mixed population of secondary structure. In the model, a mixed alpha-beta structure is observed along with nearly 35% unstructured regions. In **Figure 5A**, the simulations of wing domain in isolation up to 500 ns is shown. The radius of gyration, RMSD and RMSF values have been plotted after 500 ns of simulations (**Figure 5A-5D**). The RMSD values reach a plateau phase above 0.7 nm which indicates a significant level of structural dynamics (**Figure 5C)**. From the RMSF values, these fluctuations are mostly contributed by the residues in the long flexible loop with residues 90-130 (**Figure 5D)**. This region, as mentioned above, harbours important residues for interaction with membranes and antibodies. It has also been speculated that this may be involved in stabilising the hexamer formation. Further, the region around the greasy finger residues, which is an important part of the hydrophobic surface, also shows high RMSF fluctuations. This indicates that these regions of the wing domain may exhibit dynamic conformations in different environment and upon interaction with different binding partners.

**Figure 5.**
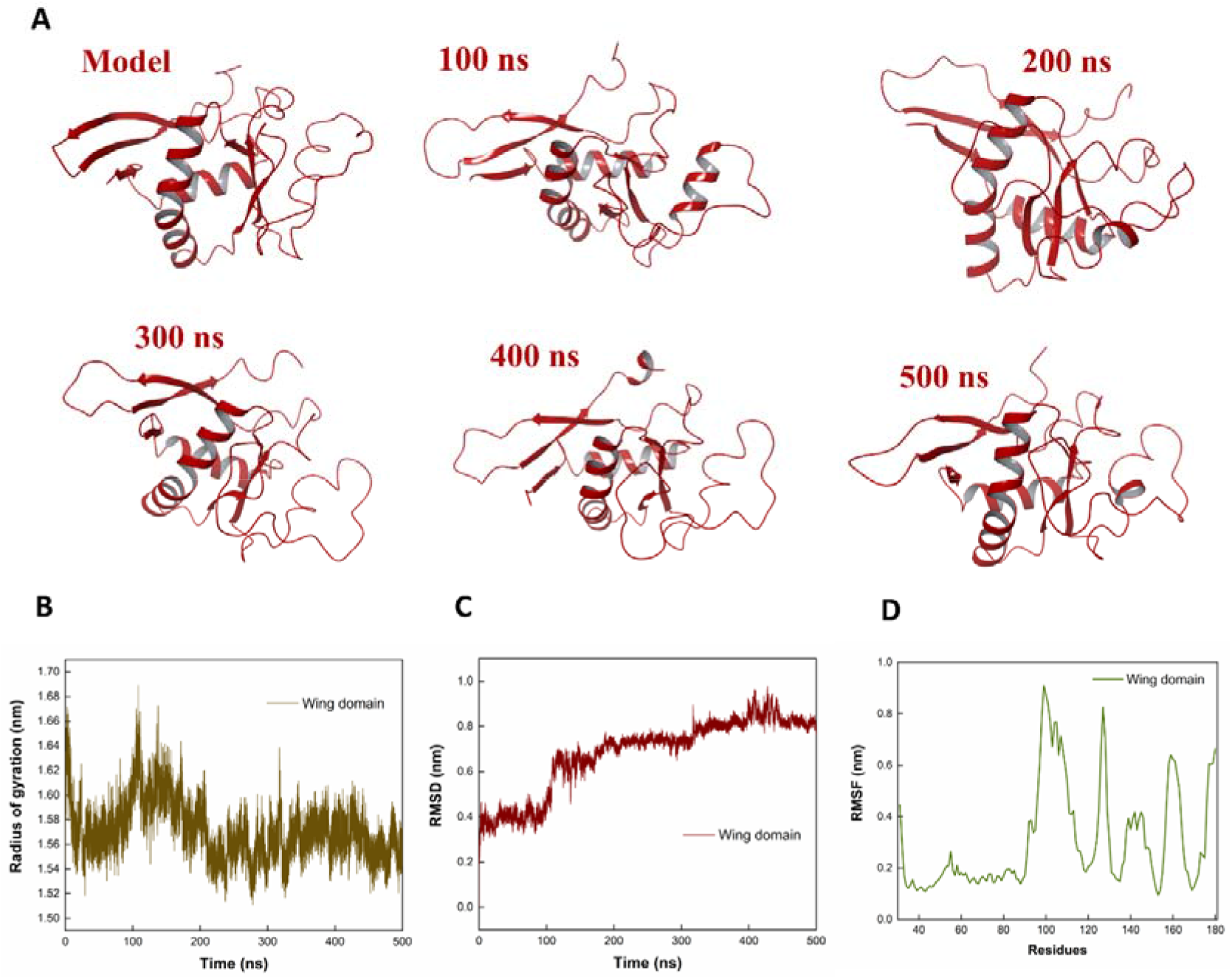
Molecular Dynamic simulations of ZIKV NS1 wing domain in isolation in water up to 500 ns (A) ZIKV NS1 wing domain in native water environment at 100, 200, 300, 400 and 500 ns. (B), (C), and (D) show the graphical representations of Radius of gyration, RMSD and RMSF, respectively.

#### 2.5 Conformational dynamics of ZIKV NS1 β-roll

##### 2.5.1 MD Simulations of ZIKV NS1 β-roll suggests it an intrinsically disordered region in isolation

The first domain towards the N-terminal of the NS1 protein is the β-roll domain. Where the β-hairpin regions of two monomers intertwine and form a mini domain swap, roll-like structure which is termed as the β-roll dimerization domain.^7^ It is a part of the discontinuous hydrophobic surface formed in the inner face of the dimer. β-roll domain is important for certain functions of NS1 such as interaction with membranes, lipids and viral proteins. In WNV NS1, it was shown that the residues 10-11 that fall in this domain may be involved in interaction with NS4B.^20^

As elucidated in the dimeric crystal structure, the beta roll domain has two beta strands in each monomer but in model, it also has a small helical region which corresponds with the secondary structure prediction made by PsiPred (**Figure S2B**). To confer the conformational dynamics, the MD simulations were performed in isolation. **Figure 6A** shows the simulations of β-roll domain in isolation up to 500 ns. The radius of gyration, RMSD and RMSF values have been plotted after 500 ns of simulations. From the simulations data, it is observed that the β-roll domain in isolation is showing a high structural instability and a propensity towards disorder. The structure became completely extended at 100 ns, 200 ns and 500 ns of simulations. RG and RMSD values have continuously showed high levels of fluctuations on simulations during entire timescale of 500 ns **(Figure 6B & 6C)**. The RMSF values over the entire length of the peptide, stay above 0.6 nm **(Figure 6D)**. This indicates a high propensity of dynamic conformational flexibility of the β-roll domain in isolation.

**Figure 6.**
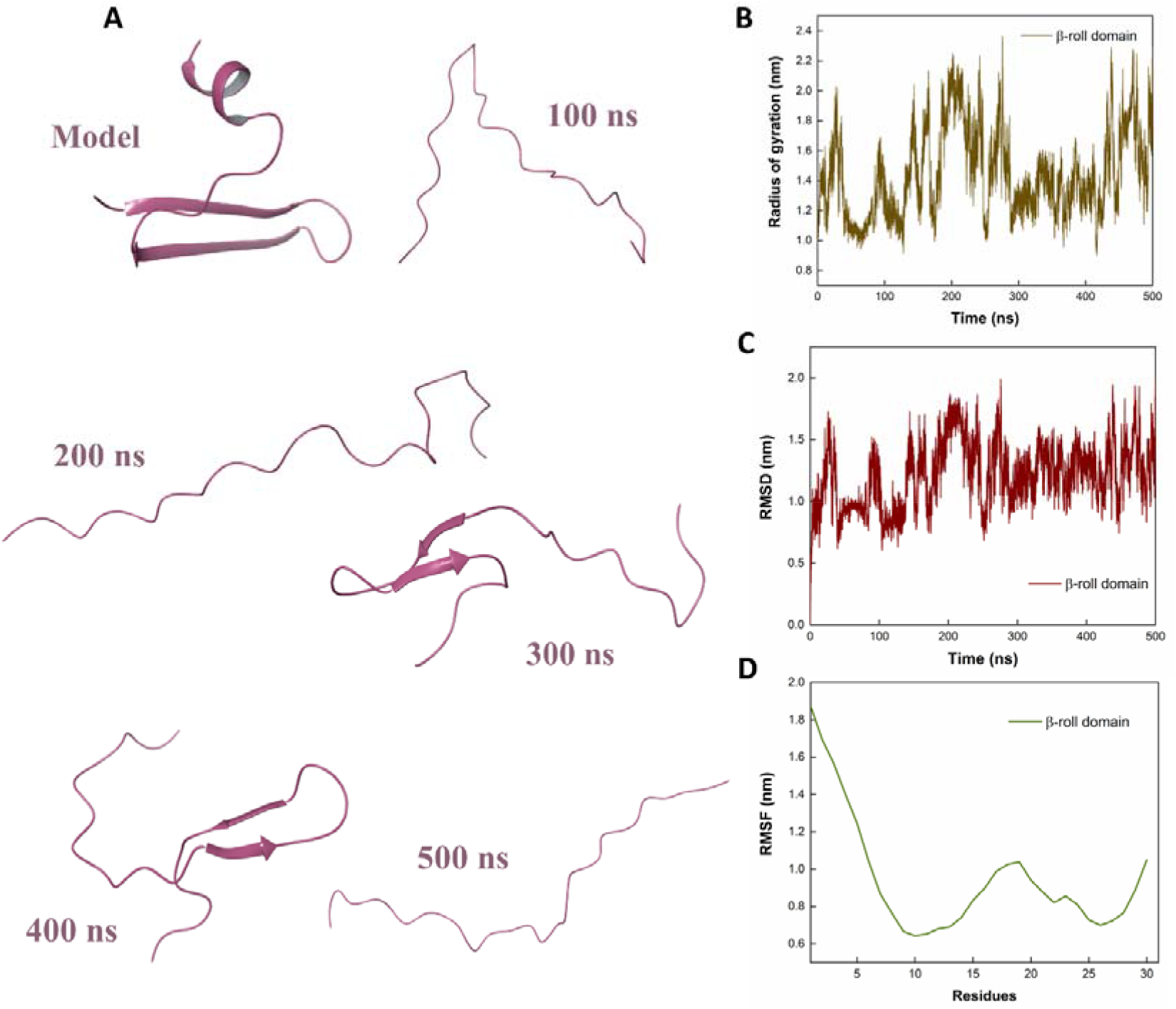
Molecular Dynamic simulations of ZIKV NS1 β-roll domain in isolation (A) Trajectory snapshots in aqueous environment at 100, 200, 300, 400 and 500 ns. (B), (C), and (D) show the graphical representations of Radius of gyration, RMSD and RMSF, respectively.

##### 2.5.2 ZIKV NS1 β-roll domain is intrinsically disordered in isolation

The CD spectra of ZIKV NS1 β-roll domain in isolation at concentration of 40 μM shows a negative peak between 198 nm - 200 nm **(Figure 7)**, which is a characteristic of unstructured/disordered proteins.

**Figure 7.**
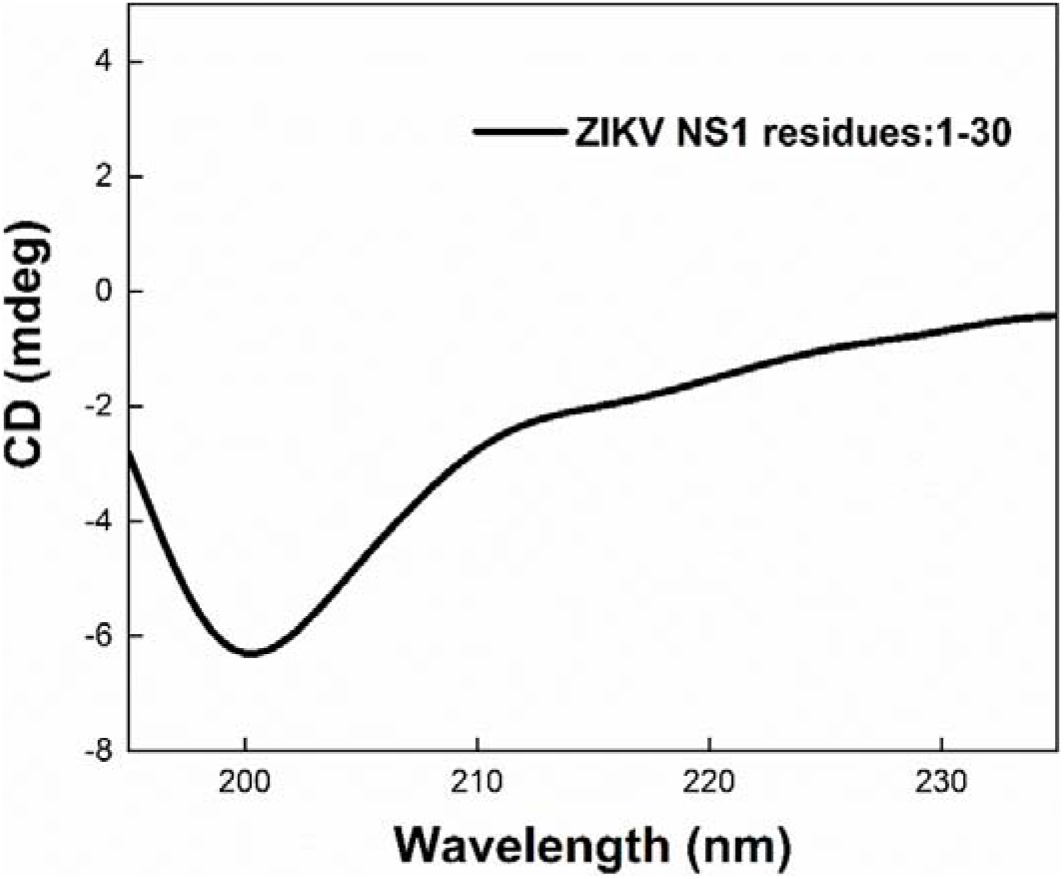
The synthesised peptide of ZIKV NS1 β-roll domain gives a disordered CD spectrum in isolation.

##### 2.5.3 Conformational changes in ZIKV NS1 β -roll domain under different lipid environment

As covered in the introduction, NS1 is capable of interacting with various cellular factors. By virtue of its β-roll domain, it even interacts with cell membranes and lipid rafts in the form of dimer and hexamer. The effect of SDS, which forms a membrane mimicking micellar structure and DOPS, DOPC, and DOTAP which were used to prepare LUVs was studied using CD spectroscopy **(Figure 8)** on the structure of ZIKV NS1 β-roll domain. The SDS micelles are used to prepare an amphipathic membrane that mimics the interface between hydrophobic and hydrophilic environments to study protein-membrane interaction.^21^ DOPC, DOPS and DOTAP were used to construct neutral, anionic and cationic LUVs, respectively.

**Figure 8.**
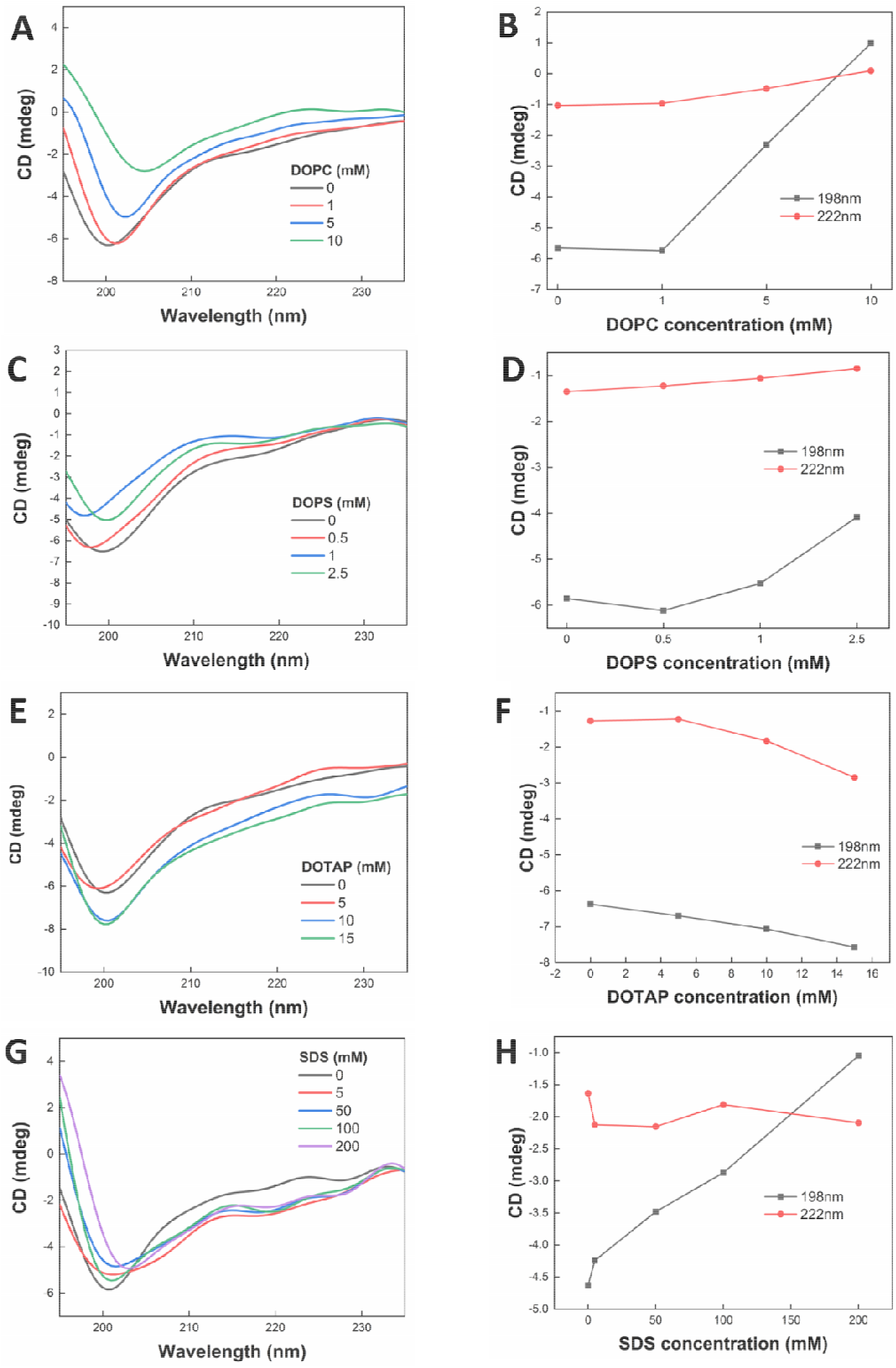
CD spectra of ZIKV NS1 β-roll domain under varying concentration of DOPC, DOPS, DOTAP, and SDS in (A), (C), (E) and (G), respectively, show conformational changes. (B), (D), (F), and (H) show the changes in ellipticity at 198 nm and 222 nm to analyze conformational transition of CD spectra.

Any significant structural changes are not observed in the lipid mimetic environments provide by DOPC (**Figure 8A and 8B**), DOPS (**Figure 8C and 8D**), DOTAP (**Figure 8E and 8F**) and SDS (**Figure 8G and 8H**). This shows that the β-roll domain of ZIKV NS1, in isolation, is not susceptible to gain-of-structure in provided lipid mimetic environment.

##### 2.5.4 Conformational dynamics in the presence of TFE

The 2,2,2-trifluoroethanol (TFE) is a strong alpha helix inducer and this is used to see the possibility of alpha helix forming propensity of peptides and proteins.^22,23^ TFE strengthens the intra H-bonds of an α-helix and weakens the H-bonding of water with the CO and NH groups of the protein.^22,23^ It has also been speculated that TFE structures the solvent in such a way that it destabilises the unfolded conformations leading to an elevated folded population.^22,23^

A significant increase in α-helical content is observed in the presence of TFE (**Figure 9A and 9B**). However, a complete transition is not seen, rather, a molten globule like conformation is observed. This behaviour was further confirmed by fluorescence lifetime measurement. Any changes in fluorophore physicochemical environments lead to the change in its property and, consequently, the fluorescence lifetime (**Figure 9C and Table 1**). Our findings reveal interactions with TFE leading to significant changes in the secondary structure.

**Table 1:** Average lifetime and chi-square values for ZIKV NS1 B-roll domain in different concentrations of TFE. The concentration of protein was kept constant at 40 _μ_**M**.

| Percentage TFE | Average lifetime (ns) | Chi-square |
| --- | --- | --- |
| 0 | 2.11 | 1.12 |
| 20 | 2.22 | 1.19 |
| 35 | 2.00 | 1.23 |
| 70 | 1.72 | 1.17 |

**Figure 9.**
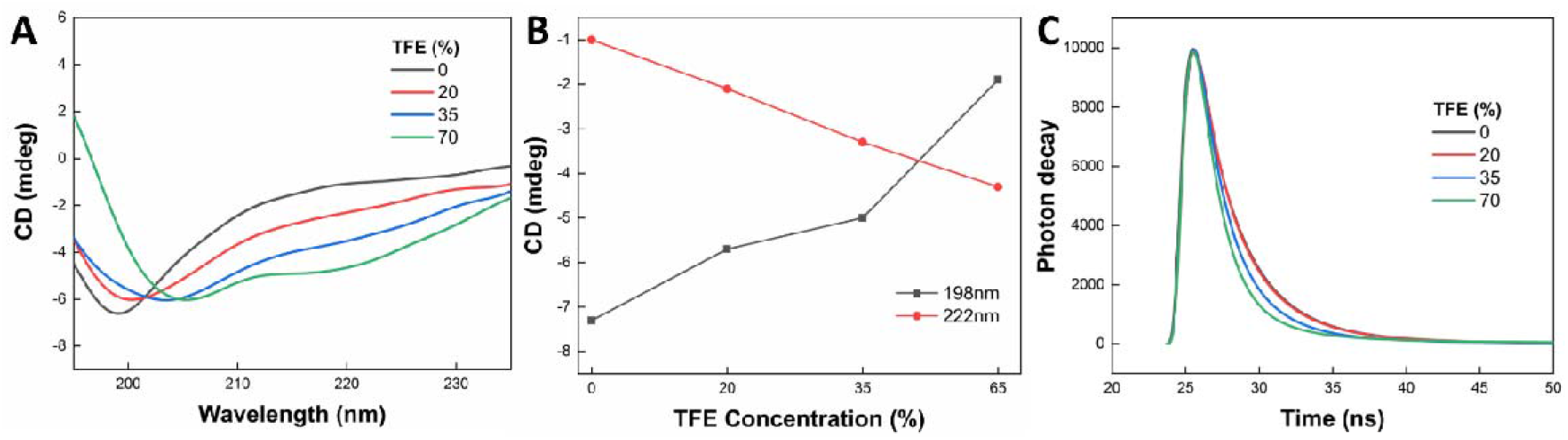
(A) CD spectra of ZIKV NS1 β-roll domain under varying concentration of TFE show conformational changes. (B) Shows the changes in ellipticity at 198 nm and 222 nm to analyze conformational transition of CD spectra. (C) Fluorescence lifetime decay curve of ZIKV NS1 β-roll domain in various concentrations of TFE.

##### 2.5.5 MD Simulations of ZIKV NS1 β-roll in lipid mimetic environment

Through our experimental approaches we observed that the ZIKV NS1 β-roll domain in isolation doesn’t show significant structural changes in the presence of lipid mimetic environment. Therefore, we performed MD simulations of ZIKV NS1 β-roll domain, in isolation, in the presence of lipids, DOPC and DOPS to confirm the folding behaviour of the peptide that we observed in the experiments. The ZIKV NS1 β-roll domain model was prepared as described earlier. It was simulated using GROMACS software up to 100 ns in the presence of 220 molecules of DOPC (**Figure 10**) and DOPS (**Figure 11**), separately. The structural dynamics were analysed using the Radius of Gyration, RMSD and RMSF.

**Figure 10.**
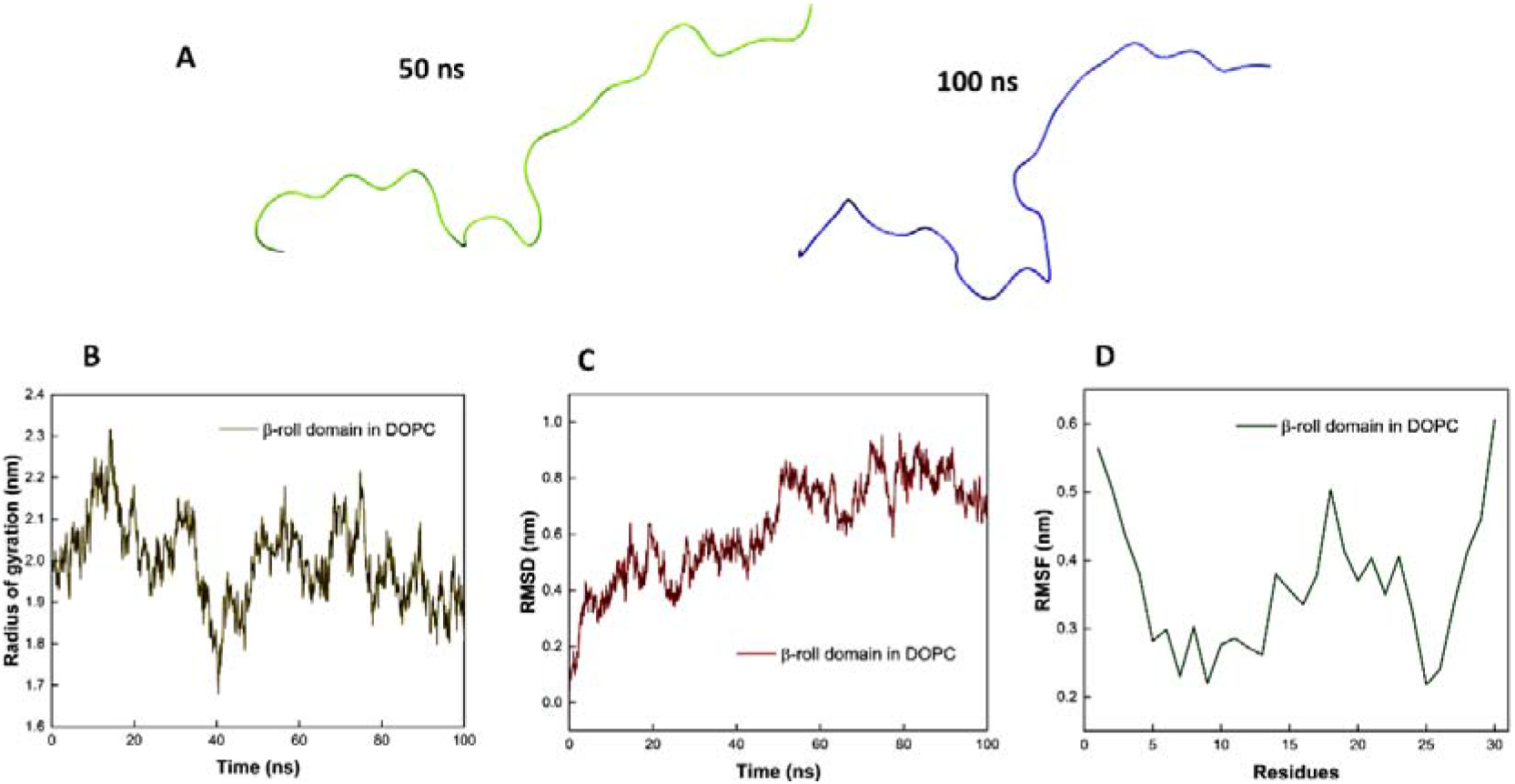
(A) Trajectory snapshots of ZIKV NS1 β-roll domain in the presence of DOPC, at 50 ns and 100 ns of MD simulations. (B), (C), and (D) show the graphical representations of Radius of gyration, RMSD and RMSF, respectively, on simulation of ZIKV NS1 β-roll domain in isolation in DOPC environment on simulations up to 100 ns.

**Figure 11.**
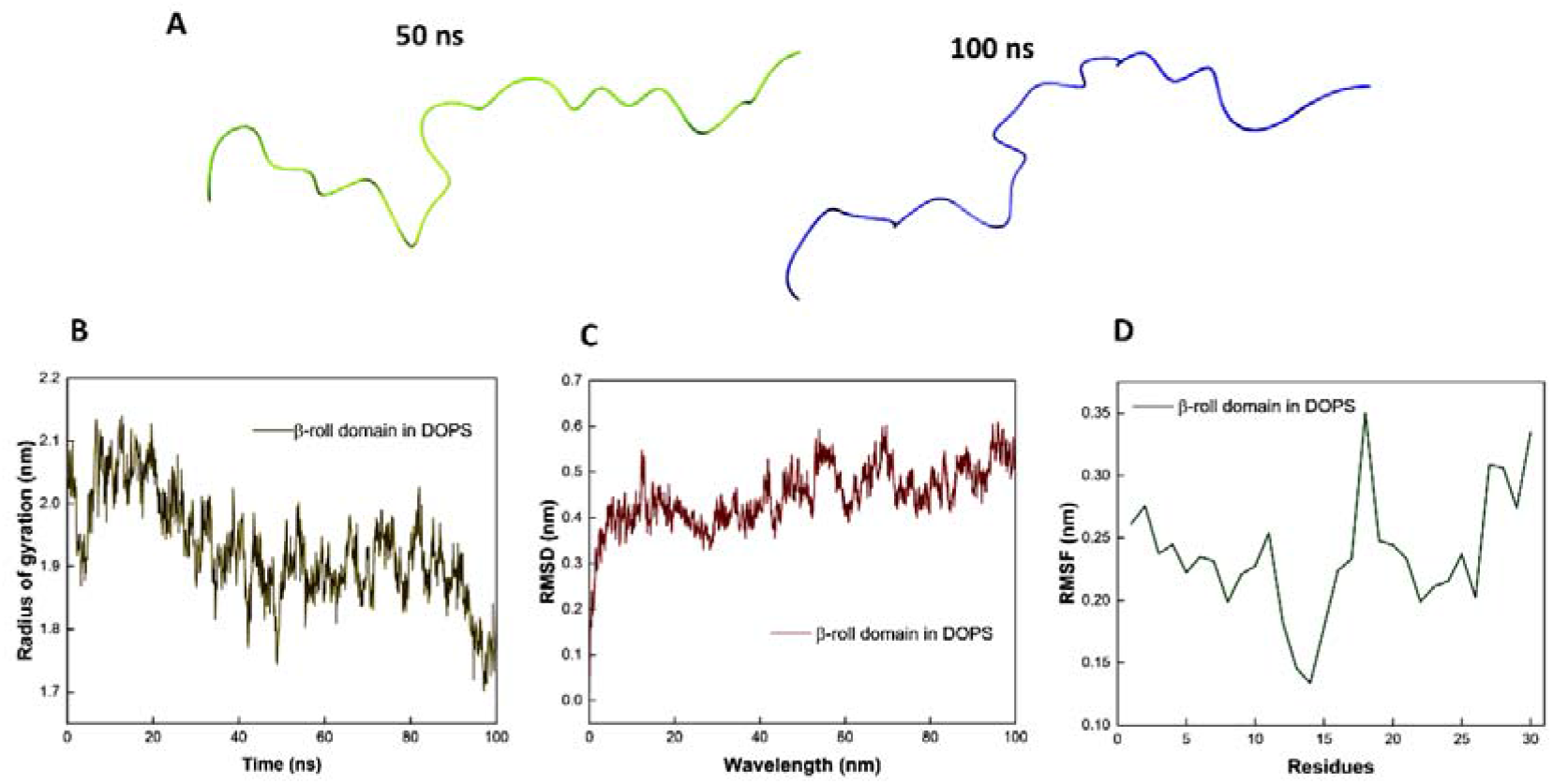
(A) Trajectory snapshots of ZIKV NS1 β-roll domain in the presence of DOPS, at 50 ns and 100 ns of MD simulations. (B), (C), and (D) show the graphical representations of Radius of gyration, RMSD and RMSF, respectively, on simulation of ZIKV NS1 β-roll domain in isolation in DOPS environment on simulations up to 100 ns.

**Figure 12:**
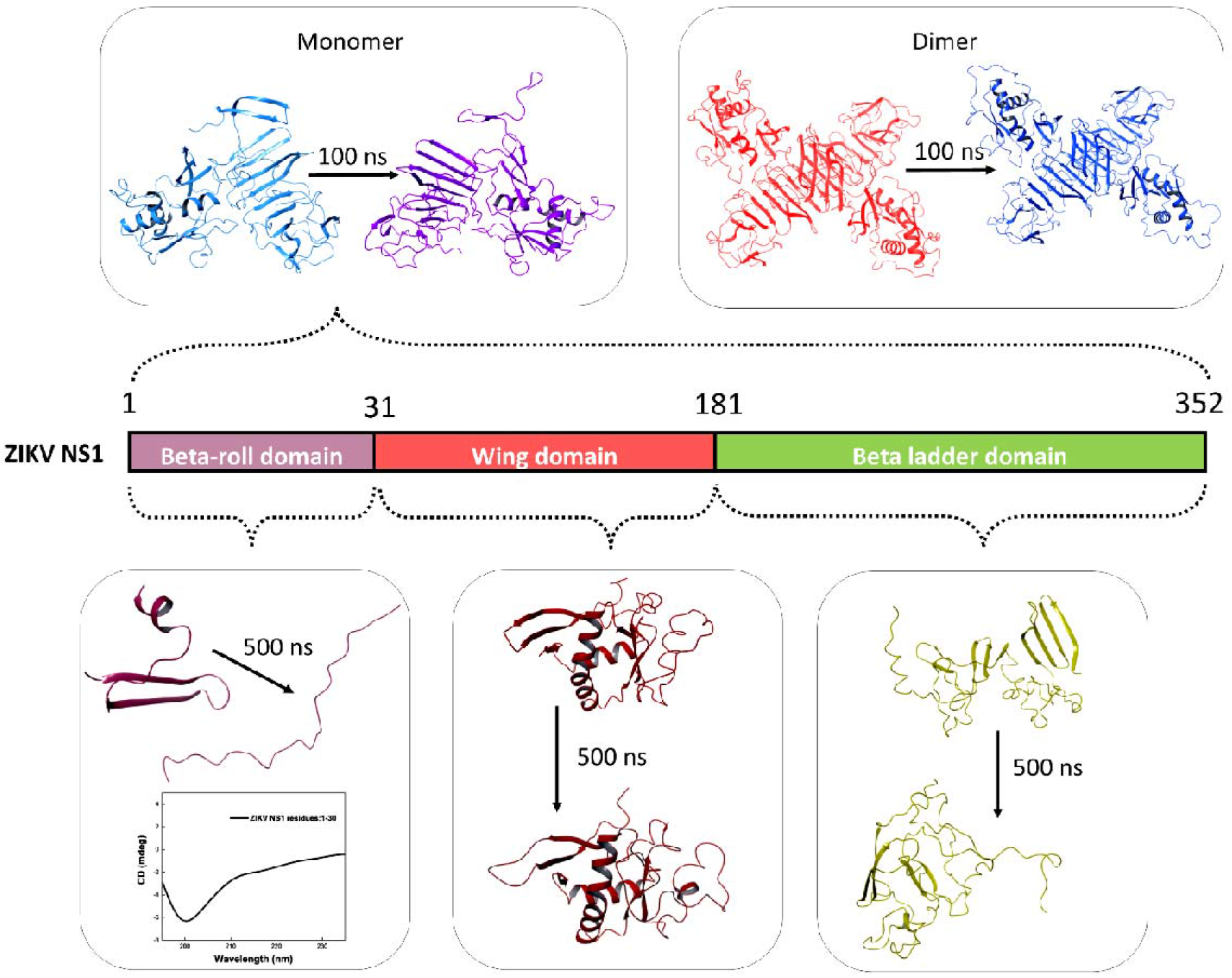
The proposed models and the MD simulations performed in this study to understand the conformational dynamics of ZIKV NS1 protein with a special focus on the β-roll domain.

DOPC was used to construct neutral liposomes and DOPS was used to construct anionic liposomes. Neutral lipids are commonly present on the outer layer of cell membranes. Golgi complex and ER membranes also contain a significant amount of neutral lipids. Anionic lipids are commonly found on the cytosolic layers of the Golgi and ER membranes, which are involved in intracellular secretory vesicle trafficking.^24,25^ The peptide model has maintained its extended structure throughout the simulations as observed from the Radius of Gyration, RMSD and RMSF plots which show high levels of fluctuations. This confirms the experimental results where we observed that ZIKV NS1 β-roll domain in isolation is resistant to gain-of-structure in lipid mimetic environment.

NS1 monomer has six disulphide bonds within the monomer itself. So, one would expect the monomer to be a pretty rigid structure. However, significant fluctuations were observed during the simulations of the full length monomer, at the β-roll domain and the wing domain. These two regions of the protein have also been implicated in interactions with various cellular factors. Further, there is a possibility that β-roll domain attains its structure when present as a dimer and is aided by other regions of the protein that are brought together spatially. Two connector regions that form a sub-domain consisting of amino acids 30 – 37 and 152 –180 that form a discontinuous sub domain, pack against the β-roll domain and connects the wing with the B-ladder domain.^7^ The β-roll and connector subdomain form a protrusion that together are responsible for the hydrophobic face of the dimer. This hydrophobic nature is conserved across flaviviruses. It has been speculated that this entire hydrophobic region helps in stabilizing the viral replication complex formation via interaction with viral NS4A/NS4B proteins.^26^ Further the greasy finger domain is a part of the connector sub-domain. Mutations in this region were deleterious to viral replication.^27^

Recently, a report has shown how the structural flexibility of β-roll domain of DENV NS1 may determine the loose teramer or stable tetramer formation of the secretory NS1.^28^ As per our observations, β-roll domain of ZIKV is disordered in isolation. Significant structural changes in the presence of membrane mimetics and membrane co-solvent mimetics is not observed. Therefore, we confirmed this behavior using molecular dynamics simulations which showed the β -roll domain to be disordered. Even though this behavior is contrary to the reported crystal structure of the NS1 dimer, it corresponds well with our simulations of full length NS1 monomer, **Figure 3B**. We observe that the β-roll domain (in full length monomer) loses its structure and becomes extended after simulation at 100ns in native water environment. As described earlier, on simulation of the dimer up to 100 ns, the RMSF values of both the chains of the dimer do not show significant fluctuations at N-terminal region, corresponding to the β-roll domain, contrary to what was observed in the simulations of the monomer. This may indicate that the interaction of the β-roll domain, which is involved in oligomerization, with the β-roll domain of the other chain, restricts the conformational flexibility of this region.

## Conclusion

In short, since ZIKV NS1 β-roll domain in isolation remains unstructured in water and even in membrane mimicking lipids and co-solvents. This suggests a possibility that the protein needs to dimerize to be functional upon lipid and membrane interaction. Our unbiased study has revealed lose core in the monomer form which are stabilized in the dimer form. Our approach was to investigate and compare the flexibility and dynamics of different domains in isolation to reveal the range of possible dynamics. Further experimental studies need to be done to explore NS1 conformational dynamics in monomeric form. Seeing how the conformational rigidity is important for the functional form of NS1, i.e., the dimer, it may be a good strategy to target the monomer to prevent the dimer formation and inhibitor discovery thereon.

## Author Contributions

RG: Conception, design, and review of the manuscript. SKK and PK: acquisition and interpretation of data and writing of the manuscript.

## Acknowledgements

All the authors would like to thank IIT Mandi and HPC utility for providing facilities. RG would like to acknowledge the Science and Engineering Research Board (SERB) (CRG/2019/005603), Department of Biotechnology, (BT/11/IYBA/2018/06), MHRD-SPARC (SPARC/2018-2019/P37/SL), and Indian Council of Medical Research (58/6/2020/PHA/BMS, and 52/04/2020/BIO/BMS). SKK is grateful to the ICMR for her senior research fellowship for Funding.

## References

1. Chibueze, E. C. et al. Zika virus infection in pregnancy: a systematic review of disease course and complications. Reprod. Health 14, 28 (2017).

2. Rawal, G., Yadav, S. & Kumar, R. Zika virus: An overview. J. Fam. Med. Prim. Care 5, 523 (2016).

3. Wang, A., Thurmond, S., Islas, L., Hui, K. & Hai, R. Zika virus genome biology and molecular pathogenesis. Emerg. Microbes Infect. 6, (2017).

4. r, R. R. & Je, L. The Dengue Virus Nonstructural Protein 1 (NS1) Is Secreted from Mosquito Cells in Association with the Intracellular Cholesterol Transporter Chaperone Caveolin Complex. J. Virol. 93, (2019).

5. Lindenbach, B. D. & Rice, C. M. Genetic interaction of flavivirus nonstructural proteins NS1 and NS4A as a determinant of replicase function. J. Virol. 73, 4611– 4621 (1999).

6. Akey, D. L., Brown, W. C., Jose, J., Kuhn, R. J. & Smith, J. L. Structure-guided insights on the role of NS1 in flavivirus infection. Bioessays 37, 489–494 (2015).

7. Brown, W. C. et al. Extended surface for membrane association in Zika virus NS1 structure. Nat. Struct. Mol. Biol. 23, 865–867 (2016).

8. Glasner, D. R., Puerta-Guardo, H., Beatty, P. R. & Harris, E. The Good, the Bad, and the Shocking: The Multiple Roles of Dengue Virus Nonstructural Protein 1 in Protection and Pathogenesis. Annu. Rev. Virol. 5, 227 (2018).

9. Gutsche, I. et al. Secreted dengue virus nonstructural protein NS1 is an atypical barrel-shaped high-density lipoprotein. Proc. Natl. Acad. Sci. U. S. A. 108, 8003–8008 (2011).

10. Edeling, M. A., Diamond, M. S. & Fremont, D. H. Structural basis of flavivirus NS1 assembly and antibody recognition. Proc. Natl. Acad. Sci. U. S. A. 111, 4285–4290 (2014).

11. Soto-Acosta, R. et al. The increase in cholesterol levels at early stages after dengue virus infection correlates with an augment in LDL particle uptake and HMG-CoA reductase activity. Virology 442, 132–147 (2013).

12. Kumar, A., Kumar, P. & Giri, R. Zika virus NS4A cytosolic region (residues 1-48) is an intrinsically disordered domain and folds upon binding to lipids. Virology 550, 27– 36 (2020).

13. Saumya, K. U., Kumar, D., Kumar, P. & Giri, R. Unlike dengue virus, the conserved 14-23 residues in N-terminal region of Zika virus capsid is not involved in lipid interactions. Biochim. Biophys. acta. Biomembr. 1862, (2020).

14. Kumar, P. et al. Reprofiling of approved drugs against SARS-CoV-2 main protease: an in-silico study. J. Biomol. Struct. Dyn. (2020) doi:10.1080/07391102.2020.1845976.

15. Giri, R., Kumar, D., Sharma, N. & Uversky, V. N. Intrinsically Disordered Side of the Zika Virus Proteome. Front. Cell. Infect. Microbiol. 6, (2016).

16. Gonçalves, R. L. et al. Dynamic behavior of Dengue and Zika viruses NS1 protein reveals monomer-monomer interaction mechanisms and insights to rational drug design. J. Biomol. Struct. Dyn. 38, 4353–4363 (2020).

17. Poveda-Cuevas, S. A., Barroso Da Silva, F.L. & Etchebest, C. How the Strain Origin of Zika Virus NS1 Protein Impacts Its Dynamics and Implications to Their Differential Virulence. J. Chem. Inf. Model. 61, 1516–1530 (2021).

18. Xu, X. et al. Contribution of intertwined loop to membrane association revealed by Zika virus full-length NS1 structure. EMBO J. 35, 2170–2178 (2016).

19. Scaturro, P., Cortese, M., Chatel-Chaix, L., Fischl, W. & Bartenschlager, R. Dengue Virus Non-structural Protein 1 Modulates Infectious Particle Production via Interaction with the Structural Proteins. PLoS Pathog. 11, (2015).

20. Youn, S. et al. Evidence for a genetic and physical interaction between nonstructural proteins NS1 and NS4B that modulates replication of West Nile virus. J. Virol. 86, 7360–7371 (2012).

21. Tulumello, D. V. & Deber, C. M. SDS micelles as a membrane-mimetic environment for transmembrane segments. Biochemistry 48, 12096–12103 (2009).

22. Roccatano, D., Colombo, G., Fioroni, M. & Mark, A. E. Mechanism by which 2,2,2-trifluoroethanol/water mixtures stabilize secondary-structure formation in peptides: a molecular dynamics study. Proc. Natl. Acad. Sci. U. S. A. 99, 12179–12184 (2002).

23. Luo, P. & Baldwin, R. L. Mechanism of helix induction by trifluoroethanol: a framework for extrapolating the helix-forming properties of peptides from trifluoroethanol/water mixtures back to water. Biochemistry 36, 8413–8421 (1997).

24. Van Meer, G., Voelker, D. R. & Feigenson, G. W. Membrane lipids: where they are and how they behave. Nat. Rev. Mol. Cell Biol. 9, 112–124 (2008).

25. Casares, D., Escribá, P.V. & Rosselló, C. A. Membrane Lipid Composition: Effect on Membrane and Organelle Structure, Function and Compartmentalization and Therapeutic Avenues. Int. J. Mol. Sci. 20, (2019).

26. Płaszczyca, A. et al. A novel interaction between dengue virus nonstructural protein 1 and the NS4A-2K-4B precursor is required for viral RNA replication but not for formation of the membranous replication organelle. PLoS Pathog. 15, (2019).

27. Akey, D. L. et al. Flavivirus NS1 structures reveal surfaces for associations with membranes and the immune system. Science 343, 881–885 (2014).

28. Shu, B. et al. CryoEM structures of the multimeric secreted NS1, a major factor for dengue hemorrhagic fever. bioRxiv 2022.04.04.487075 (2022) doi:10.1101/2022.04.04.487075.

