## Supplementary file for "Structural dynamics of Zika Virus NS1 via a reductionist approach reveal the disordered nature of its beta roll domain in isolation"

1. **Mass spectrum and HPLC spectrum of synthesized peptide**

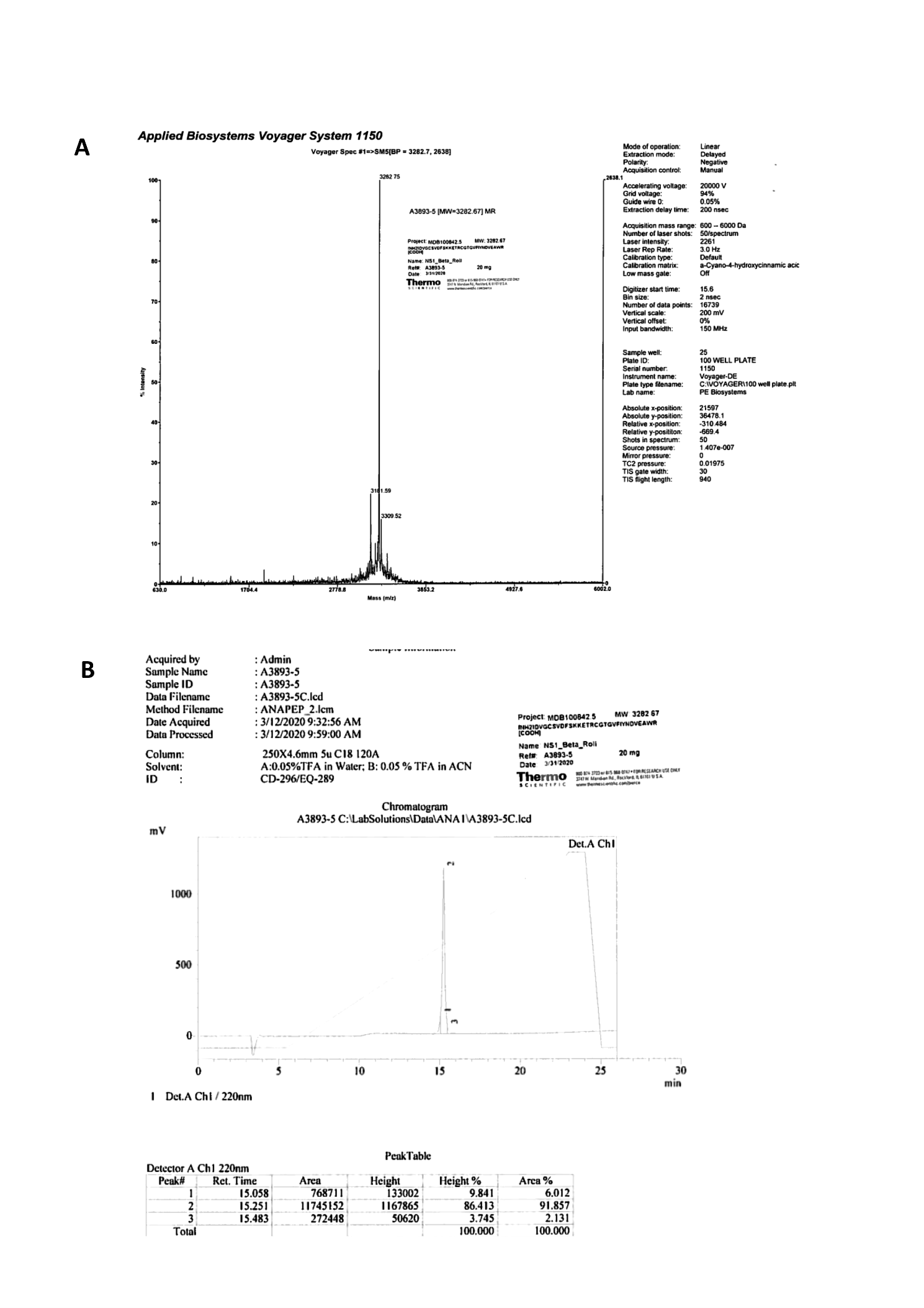

**Figure S1:** (A) Mass spectrum and (B) HPLC spectrum of synthesized peptide.

2. **The secondary structure and disorder propensity prediction of ZIKV NS1 and its domains in isolation.**

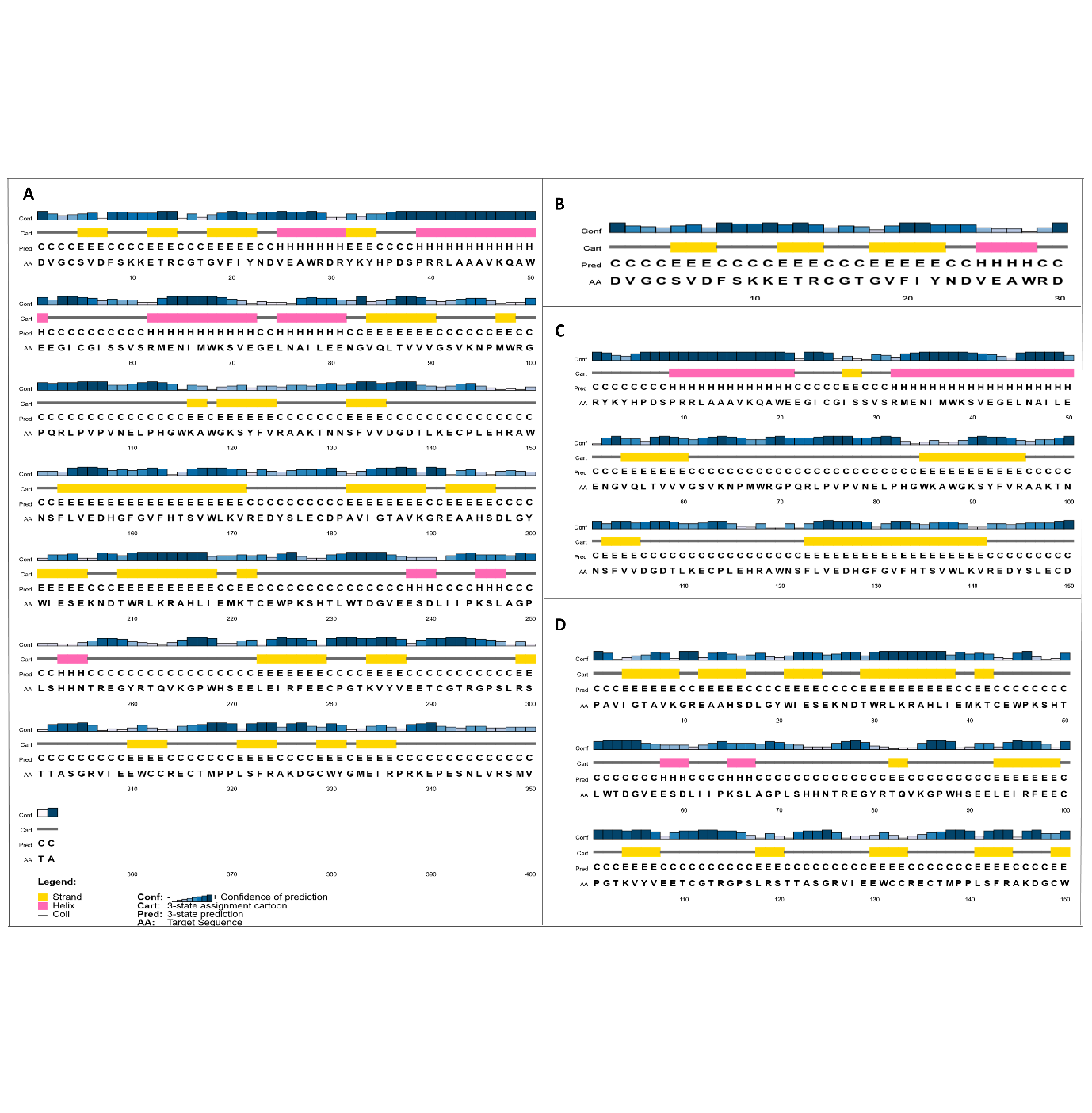

**Figure S2:** Secondary structure prediction of A) ZIKV NS1 and its domains in isolation B) β-roll domain, C) wing domain and D) ladder domain by PsiPred. (Helix, beta strand and random coil are represented by pink, yellow and grey, respectively.)
